# Florigen governs shoot regeneration

**DOI:** 10.1101/2021.03.16.435564

**Authors:** Yaarit Kutsher, Michal Fisler, Adi Faigenboim, Moshe Reuveni

## Abstract

It is widely known that during the reproductive stage (flowering), plants do not root well. Most protocols of shoot regeneration in plants utilize juvenile tissue. Adding these two realities together encouraged us to studied the role of florigen in shoot regeneration. Mature tobacco tissue that expresses the endogenous tobacco florigen mRNA regenerates poorly, while juvenile tissue that does not express the florigen regenerates shoots well. Inhibition of Nitric Oxide (NO) synthesis reduced shoot regeneration as well as promoted flowering and increased tobacco florigen level. In contrast, the addition of NO (by way of NO donor) to the tissue increased regeneration, delayed flowering, reduced tobacco florigen mRNA. Ectopic expression of florigen genes in tobacco or tomato decreased regeneration capacity significantly. Overexpression pear *PcFT2* gene increased regeneration capacity. During regeneration, florigen mRNA was not changed. We conclude that florigen presence in mature tobacco leaves reduces roots and shoots regeneration and is the possible reason for the age-related decrease in regeneration capacity.

## Introduction

Plant regeneration by rebuilding new organs (organogenesis) results from new organ formation through dedifferentiation of differentiated plant cells and reorganization of cell division to create new organ meristems and new vascular connection between the explant and the newly regenerating organ ^1,2^. Regeneration of a multicellular organism from a segment of adult somatic tissue is a prevalent phenomenon in both plants and animals^3^. While plant cells are presumed to maintain totipotency, animal cells, except for stem cells, do not. Plant cells that are already differentiated into organs can regenerate whole plants under *in vitro* conditions^4,5^. The ratio of the two main growth regulators, auxin and cytokinin, determines regenerating cells’ developmental program. High cytokinin to auxin ratio directs shoot regeneration, while low cytokinin to auxin ratio results in roots^6^.

Animals contain stem cells that maintain totipotency throughout the animal’s life for the purpose of tissue regeneration and repair. In contrast, plants, except for two cells at the apical meristem and two cells at the root meristem, do not have stem cells or totipotent cells circulating. Interestingly but in common with animal stem cells that decline in regenerative ability with age, there is a progressive decline in plant cells’ regenerative ability during their life cycle^7,8^. In plants, chronological age affects the plant regenerative capability manifested as rooting ability or shoot regeneration ^9,10^. Recently Zhang et al.^11^ showed that *miR156*, a chronologically regulated microRNA, regulates shoot regeneration in leaf segments from tobacco plants by allowing the gradual changes in *SQUAMOSA PROMOTER BINDING PROTEIN-LIKE (SPL)* mRNA levels^11^. The rise in *SPL* mRNA levels leads to a progressive drop in the ability to regenerate shoots in tobacco and *Arabidopsis* explants^11^. *miR156*-targeted *SPLs* were shown to regulate a diverse age-related process, such as embryonic pattern formation, juvenile-to-adult phase transition, the timing of flowering, and shoot regeneration ability^11,12,13,14,15,16,17^. Thus, the juvenile to adult change is strictly controlled by a regulatory gene network that contains at least *miR156* that regulated *SPL* family members and downstream of *SPL* like *FLOWERING LOCUS T* (*FT*)^18^.

Research reports in past years have shown that flower buds’ presence reduces rooting, such as in Rhododendron and *Camellia, Coleus, Vaccinium*, and *Taxus*^19^. Even earlier, Wilton^20^ showed that little or no cambial activity is present in the stems of flowering plants compared to non-flowering plants^20^. The above effects indicate that the capacity to regenerate roots or shoots and maintain cell division activity in the cambium is associated with the plants’ maturity state. It is a common practice in vegetative rooting of cuttings to use as juvenile tissue as possible.

*FLOWERING LOCUS T (FT)* is a transmissible floral inducer in flowering plants. *FT* is a critical element in annual plants’ competence to flower shortly after emergence; however, perennial plants contain at least two *FLOWERING LOCUS T* genes with different functions^21^. Perennials have an extended juvenile vegetative period lasting up to many years of vegetative growth before achieving flowering^18,22^. After flowering, perennials enter into a yearly cycle of vegetative and reproductive processes. Perennials express at least two versions of *FT*-like genes to regulate flowering and non-flowering phases. *FT1*-like and *FT2*-like (a phosphatidylethanolamine-binding protein family) of perennial plants regulate cellular proliferation and new tissue formation and induce flowering when expressed tobacco or *Arabidopsis*^21,22^. However, while *FT1*-like expression in perennial plants precedes flowering, suggesting it functions as florigen, *FT2*-like genes are associated with the juvenile and vegetative period^21,23^. Thus, *FLOWERING LOCUS T*-like genes coordinate the repeated cycles of vegetative and reproductive growth in perennial like poplar and pear by cycled expression year-round^21,24^.

Kumar et al.^25^ demonstrated that Oncidium’s flowering is mediated by NO (nitric Oxide) levels, suggesting that NO controlled the phase transition and flowering process^25^. Application of sodium nitroprusside (NO donor) to *Arabidopsis vtc1* mutant caused late flowering, and expression levels of flowering-associated genes (*CO, FT*, and *LFY*) were reduced, suggesting NO signaling is vital for flowering^25,26^. As the induction to flowering or vegetation pattern relies on the balance between the expression levels of genes in the *PEBP* gene family (phosphatidylethanolamine-binding protein) like *FT1* or *FT2*^21,24,27^ mutations in these homologous genes have different consequences on flowering or vegetative growth. But it seems that these genes determine the fate of the meristem for vegetative or reproductive growth^28^.

In the study here, we present evidence to show that the florigen genes levels in tobacco or tomato influence regeneration capacity. Overexpression of pear *PcFT2* gene increased regeneration capacity while *FT1* or florigen reduces regeneration capacity. During regeneration, tobacco florigen mRNA does not change. We conclude that florigen presence in mature tobacco leaves reduces roots and shoots regeneration, and it may be the possible reason for the age-related decrease in regeneration capacity.

## Results

### Root and shoot regeneration is affected by leaf age

We tested root regeneration from leaf petioles and found that as the leaves mature and the plant approaches flowering, the number of roots regenerated declines (Fig. 1a) as well as percent regeneration (Fig. S1). Root regeneration from mature leaf segments taken from the leaf blade was much lower than juvenile leaf segments (Fig. 1b), as was percent shoot regeneration (Fig. S2a). The number of shoots regenerated from juvenile leaves (leaf number 7-8) was significantly higher than from mature leaves (leaf number 20-21) (Fig. 1c), as was percent shoot regeneration (Fig. S2b). Juvenile leaves regenerate more roots and shoots than mature leaves. The juvenile leaves’ tips are much rounded than the mature leaves (Fig. 1d); as the tobacco plant ages, the leaves turn more elongated, which can be seen as the ratio between leaf width to leaf length decreases (Fig. S3). Under our conditions in the growth room, tobacco plants flowers at about 20 to 22 leaves.

**Fig. 1:**
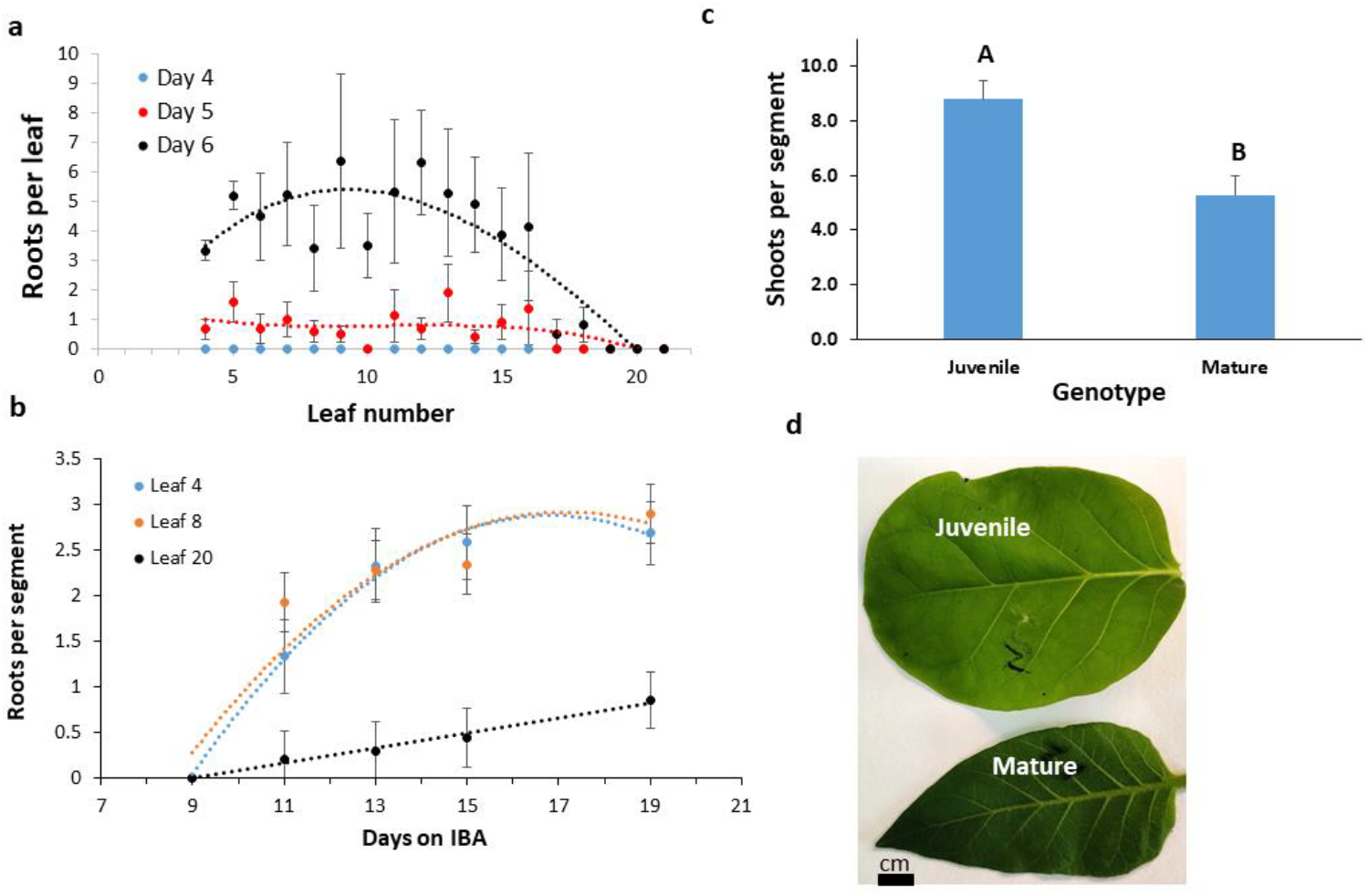
Effect of leaf age on regeneration. Tobacco leaves were detached at the plant’s petiole-stem junction and numbered starting from the first true leaf and were placed in agar supplemented with IBA. **a**: The number of roots ± SE regenerated from leaf petioles were counted after 4, 5, and 6 days in IBA each day. The experiment was done in 2 replicates with 5-7 leaves each leaf number. Percent regeneration can be seen in figure S2. **b**: The number of roots ± SE regenerated from leaf blades of juvenile (leaf 4 or 8) or mature (leaf 20) leaves. The experiment was done in 3 replicates and each 10-15 leaf segment. Percent regeneration can be seen in figure S3. **c**: The number of shoots ± SE regenerated from leaf blades was counted after 30 days on the regeneration medium of mature (leaf 20) or juvenile (leaf 8) segments. The experiment was done in 3 replicates, and each had 20-25 leaf segments. Percent regeneration can be seen in figure S3. **d**: Phenotype difference between mature and juvenile leaves. Statistical analysis was conducted among each color group using the JMP program using Tukey analysis. Different letters depict statistically significant differences between genotypes or treatments (p{f}<0.001).

### Shoot regeneration and flowering are affected by Nitric Oxide

The flowering of tobacco and *Arabidopsis* plants was affected by a short preincubation of the seedlings in the presence or absence of Nitric Oxide (NO) level modifiers. Growing the seedlings of tobacco (Fig. 2a) or *Arabidopsis* (Fig. 2b) on media containing the NO synthesis inhibitor DiPhenyleneiodonium (DiPhenyl) prior to planting in pots advances flowering of both plant genotypes (Fig 2a; 2b). Incubation with NO donor Molsidomine (Molsido) had a non-significant but repeated small flowering delay (Fig 2a; 2b). Shoot regeneration was enhanced in plants treated with Molsidomine (Fig 2c) and inhibited by DiPhenyl in non-transformed SR1 plants. Overexpression of avocado florigen in SR1 plants reduced shoot regeneration (Fig. 2c) while promoting flowering (Fig S2c). DiPhenyl inhibited shoot regeneration in florigen expressing and untransformed plants, but florigen expressing plants were insensitive to treatment with NO donor (Fig. 2c).

**Fig. 2:**
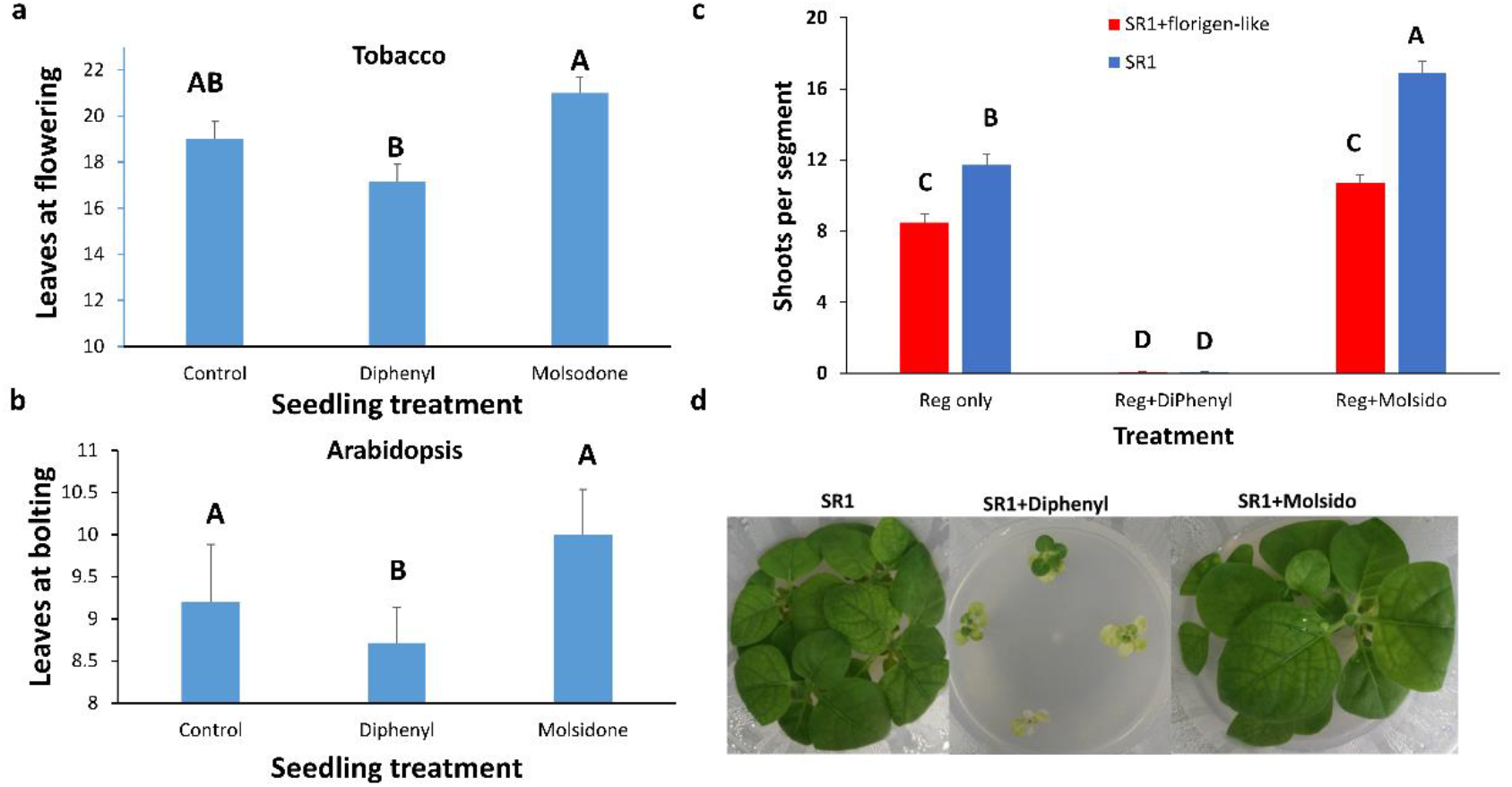
Effect of Nitric Oxide donor or inhibitor on flowering and shoot regeneration. **a**: Tobacco seedlings at the cotyledon stage were transferred and grown on an agar medium supplemented with 2% sucrose and 5μM DiPhenyl or 5μM Molsido. At the seven-leaf stage, seedlings were transferred to soil and grown to flower. The number of leaves at the time flower buds emerged ± SE was counted for tobacco plants, each treated with DiPhenyl or Molsido and control. The experiment was done in 2 replicates with 5-7 plants each. **b**: *Arabidopsis* seedlings at the cotyledon stage were grown on agar medium supplemented with 2% sucrose and 5μM DiPhenyl or 5μM Molsido. At a six-leaf stage, seedlings were transferred to soil and grown to flower. The number of leaves at the time flower buds emerged ± SE was counted, each treated with DiPhenyl or Molsido and control. **c**: The effect of Nitric Oxide donor (**Reg+Molsido**) or inhibitor (**Reg+DiPhenyl**) on the number of shoots ± SE regenerated after 30 days on juvenile leaves segments from non-transformed plants (**SR1**); plants transformed with a 35S::florigen-like gene from avocado (**SR1+florigen**), and plants transformed with a 35S::non-florigen-*PEBP*-like gene from avocado (**SR1+non-florigen**). The experiment was done in 3 replicates, and each had 20-25 leaf segments. Statistical analysis was conducted among each color group using the JMP program and Tukey analysis. Different letters depict statistically significant differences between treatments (p{f}<0.001). **d**: Tobacco seedlings treated with 5μM DiPhenyl or 5 μM Molsido or none before transfer to soil. The seedlings were transferred to 2% sucrose only or 2% sucrose ± 5μM DiPhenyl or 2% sucrose ± 5μM Molsido agar at the cotyledon stage. All plants recovered from the treatments.

Treating tobacco seedlings with DiPhenyl that inhibits NO synthesis^29,30^ shows a similar phenotype of lacking NO as in *Arabidopsis nos1* mutant^31^ (Fig. 2d) i.e., impaired growth, yellowish first true leaves, reduces root size, defective abscisic acid-induced stomatal movements, and most importantly, induces early blooming^31^. Transfer of the tobacco seedlings to the soil without DiPhenyl resulted in rapid greening and growth and early flowering (Fig. 2d). The ratio between leaf width to leaf length dud did not change after treatment with NO modifiers (Fig. S3).

### *FLOWERING LOCUS T* mRNA level is influenced by leaf position and NO treatment

The level of mRNA of several *FLOWERING LOCUS T (NtFT1, NtFT2, NtFT3*, and *NtFT4)* genes of tobacco was compared between juvenile leaf and mature leaf and juvenile leaves treated with DiPhenyl or Molsido. *NtFT4* mRNA (the tobacco florigen) level was high in mature leaves and DiPhenyl treated juvenile leaves and correlated with flowering. In leaves of plants that do not flower, *NtFT4* was not expressed (Fig 3a). *NtFT2* and *NtFT1* are expressed in all examined leaves (Fig 3a), and *NtFT3* was only expressed in mature leaves. During the initial regeneration period, the *NtFT* gene family level does not change, but *NtTFL1* expression increases (Fig. 3b).

**Fig. 3:**
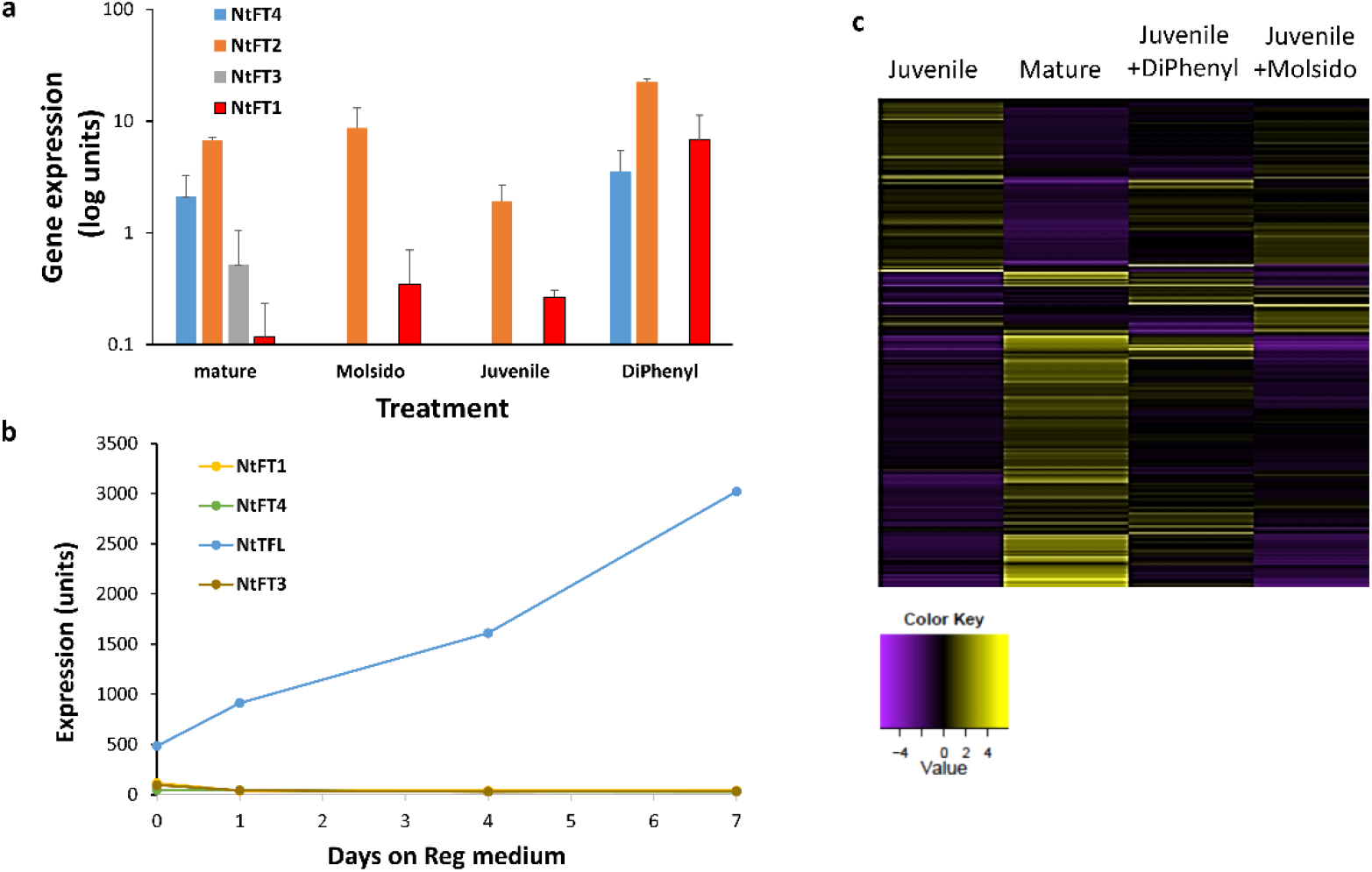
Effect of treatment with Nitric Oxide donor or inhibitor or leaf age on tobacco *FLOWERING LOCUS T* gene family expression. **a**: Tobacco seedlings at the cotyledon stage were transferred and grown on an agar medium supplemented with 2% sucrose and 5μM DiPhenyl or 5μM Molsido. mRNA levels of the *FLOWERING LOCUS T* gene family are expressed as log units. **b**: Expression level of tobacco *FLOWERING LOCUS T* gene family during the seven days induction period of shoots on Reg medium. **c**: Visualisation by heatmap of genes whose mRNA expression increase or decreases at the juvenile, mature, juvenile leaf treated with DiPhenyl and juvenile leaf treated with Molsido.

The expression of genes associated with *FLOWERING LOCUS T*, like *TEMPRANILLO* or *SQUAMOSA PROMOTER BINDING*, did not show the pattern exhibited by the tobacco florigen *(NtFT4)* (Tables S1 and S2).

mRNA expression heatmap shows extensive changes between mature and juvenile leaves (Fig 3c). The expression of 2792 genes changes between juvenile and mature leaves, while between juvenile leaves treated with DiPhenyl, only 403 genes changed expression, and between juvenile leaves treated with Molsido, only 212 genes changed expression (Fig. S4a). The gene heatmap expression of juvenile leaf treated with DiPhenyl shows specific patterns like the mature leaf and the Molsido treated leaf is similar to juvenile leaves (Fig 3c). Six different expression patterns, i.e., C1-C6, were distinguished according to similarity or difference to juvenile leaf (Fig. S4b).

### Florigen affects the leave’s ability to regenerate

Overexpression of avocado (*Presea americana*) *FLOWERING LOCUS T*-like genes (florigen) in tobacco resulted in reduced shoot regeneration capacity (Fig 4a). The flowering promoting effect of the florigen caused a decrease in shoot regeneration. In tomato cotyledon segment overexpressing either the tomato florigen (SFT) or the pepper florigen (*CaFT1*-LIKE) (Fig 4b). Mature leaf segments collected at the time flower buds are visible show reduced shoot regeneration in non-transformed plants. However, flowering did not affect shoot regeneration in plants transformed with avocado florigen (*PaFT1*) (Fig. 2c). Overexpression of pear (*PcFT2*) rejuvenator gene in tobacco plants resulted in increased shoot regeneration capacity in juvenile leaf segments (Fig 4d).

**Fig. 4:**
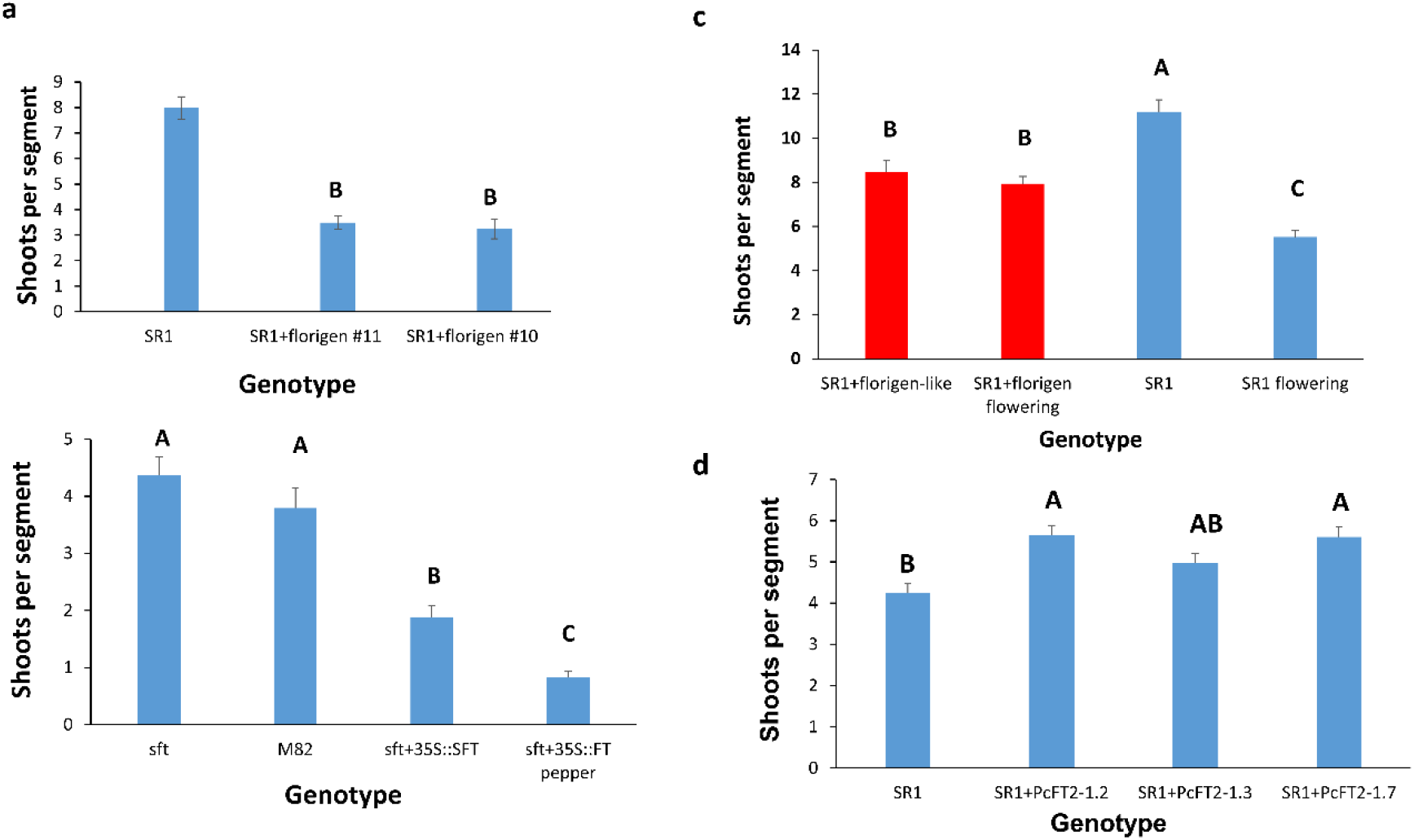
Effect florigen or rejuvenator on shoot regeneration. **a**: Number of shoots ± SE was counted for SR1 tobacco leaf segments, each transformed with florigen from avocado or non-transformed (SR1). The experiment was done in 4 replicates from separate transformed plants, each with 20 segments per plate on Reg medium. **b**: The number of shoots±SE was counted after 30 days on the regeneration medium of tomato *S. lycopersicon* cv M82. Cotyledons segments 20 per plate, four plates per genotype were tested for regeneration ability. **sft** = M82 plants mutated in florigen gene; **M82** = wild type M82 plants; **sft+35::SFT** = M82 plants mutated in tomato florigen gene overexpressing a functional tomato florigen under 35S promotor; **sft+35::FT pepper** = M82 plants mutated in tomato florigen gene overexpressing a functional *Capsicum annum* florigen (*FT1-*like) gene. The number of shoots ± SE was counted for tomato cotyledons segments placed on regeneration medium. **c**: The number of shoots ± SE was counted after 30 days on the regeneration medium of mature (noted as flowering) or juvenile tobacco leaves segments. The experiment was done in 3 replicates, and each had 20-25 leaf segments. **d**: Number of shoots ± SE was counted for SR1 tobacco leaf segments from plants overexpressing the pear (*Pyrus communis*) rejuvenator gene that confers juvenility and non-transformed tobacco (SR1). The experiment was done in 4 replicates from separate transformed plants, each with 20 segments per plate on Reg medium Statistical analysis was conducted using the JMP program and Tukey analysis. Different letters depict statistically significant differences between genotypes or treatments (p{f}<0.001).

## Discussion

We designed this study to analyze the basis of root and shoot regeneration differences between vegetative state (Juvenile) and flowering state (Mature) plant tissues. Past reports described that flower buds’ presence on a plant reduces rooting, for example, in Rhododendron, *Camellia, Coleus, Vaccinium*, and *Taxus*^19^ or inhibit cambial activity in stems of flowering plants^20^. The differences between juvenile and mature tissues in the capacity to regenerate roots or shoots depend on physiological age. Most shoot and/or root regeneration protocols vary vastly between plant species; almost with a few exceptions, use a tissue (or explant) that are juvenile, such as cotyledon, hypocotyl, petiole, or dormant meristem. In both plants and animals, regeneration ability declines with age^7,8,32^.

FLOWERING LOCUS T (FT) is a small mobile protein that functions as a floral and developmental regulator gene family. FT protein is a critical element in annual plants’ competence to flower shortly after emergence; however, perennial plants contain at least two *FT* genes with different functions in flowering florigen and a rejuvenator^21^. Perennials plants have an extended juvenile vegetative period lasting up to many years of vegetative growth before achieving flowering^18,22^. After the first flowering period, perennials enter into a yearly cycle of vegetative and reproductive processes. Perennials express at least two versions of *FT* genes florigen and a rejuvenator. Both are a phosphatidylethanolamine-binding protein family (*PEBP* gene family) and induced flowering when expressed in tobacco or *Arabidopsis*^21,22^. However, while *FT1*, the florigen expression in perennial plant’s expression, precedes flowering, the *FT2* the rejuvenator expression is during the vegetative period ^21,23^. Thus, florigen and a rejuvenator are both homologs to the *A. thaliana* gene *FLOWERING LOCUS T (AtFT1)* gene, coordinating the repeated cycles of vegetative and reproductive growth in woody perennial like poplar^24^ (Populus spp.) or pear^21^ by cycling expression year-round. Florigen action in the meristem to induce flowering^27,33^ functions to transform the leaf into a mature organ changing its shape and reducing its capacity to regenerate. This study shows that independently from flowering and other meristematic effects, *FT* functions in the leaf tissue as a determinant of juvenility or maturity depending if the rejuvenator or the florigen is expressed. Examination of FT’s immediate suspects in meristematic flowering processes did not reveal an expression pattern that follows *FT* mRNA (Table S1 and S2).

NO (nitric Oxide) levels are involved in flowering, as demonstrated in Oncidium’s^25^ and *Arabidopsis*^34^. Application of NO donor on *Arabidopsis vtc1* mutant caused late flowering, and the expression levels of flowering-associated genes *(AtCO, AtFT*, and *At*LFY) were reduced, suggesting NO signaling is part of flowering^25,26^. Reducing the amount of NO in *Arabidopsis* plants promoted flowering, increasing NO inhibited flowering^34^, as was shown here. In parallel, reducing the amount of NO decreases regeneration, and increasing the amount of cellular NO increases regeneration^30^ (and in this study). We postulated a connection between NO and *FT* gene family members; it seems that NO level controls *FT* mRNA expression and thus flowering and regeneration ability. A link between flowering and root regeneration (rooting) is known for decades and used in crop plants’ vegetative production of crop plants like trees, vegetables, and flowers. The florigen’s mRNA expression level seems to explain why plants at the reproductive stage do not regenerate as well as plants in the vegetative phase when the florigen is not expressed.

Juvenility across kingdoms is associated with enhanced regenerative ability. For example, juvenile plants exhibit a high regenerative capacity; as the plant mature, this capacity declines^32^, as shown here, and modifying mice’s adolescent state affects tissue repair, a type of regeneration^35^ or juvenile axolotl can regenerate a limb faster than an adult^36^. These observations show that the juvenility state of the tissue governs plants’ and animals’ regenerative capacity. Zhang et al.^32^ speculated that the binding of SPL9 to ARR2 changes the conformation of ARR2, thereby impairing its transcriptional activation toward downstream targets^32^. In the flowering cascade, *FT* is influenced by *miR156* and *SPL* genes^37^. Thus, as shown here, the florigen protein FT affects regeneration capacity on its own.

Our results revealed that the decrease in shoot and root regeneration in mature plant tissue is correlated with a high florigen expression. The mechanism causing the reduced shoot or root regenerative capacity in old plants and whether *FT* expression is connected to altered phytohormones response awaits further investigations. Shoot and root regeneration is influenced by many factors, the explants, the culture medium, phytohormones, and gelling agent, to name the most tested. *FT* genes expression level or presence can be used as a marker for regeneration capacity. *FT* gene manipulation can increase plant species propagation, especially in recalcitrant and rare and endangered plants.

## Materials and Methods

### Seed Sterilization and Plant Growth Conditions

Seeds of tobacco *(Nicotiana tabacum* L. cv. SR1) or *Arabidopsis* Colombia were placed in a 1.7 mL micro-tube (Eppendorf) filled with sodium hypochlorite (0.5% active material) and incubated for 5 min. The tube was shaken during the sterilization. After incubation, the seeds were rinsed three times with sterile water and spread on Petri dishes contained ½-strength MS medium (Duchefa Co. Haarlem, Netherlands, Product number M0221.0050). After about two weeks, seedlings were planted in polypropylene Vitro Vent containers (Duchefa, NL; 9.6 cm × 9.6 cm and 9 cm in height) containing the same media to obtain disinfected plants. Molsido (five μM Molsidomine and five μM DiPhenyleneiodonium chloride) was added to the agar medium when treated with NO affectors. Plants were grown in sterile boxes in a growth room with 16 hours of light and eight hours of darkness at 26 °C for weeks until leaves were ready to be harvested. Tobacco plants were transformed as described before ^30,38^ with Avocado (*Persea Americana*), FLOWERING LOCUS T-like plasmids obtained from Ziv et al.^39^, pear (*Pyrus communis*) from Frieman et al.^21^, and pepper (*Capsicum annum*) from Borovsky et al^40^. Transgenic tomato (*Solanum lycopersicon* M82) seeds were obtained from^40^.

The effect of NO on regeneration was done by placing leaf segments on Reg medium containing NO donors (5 μM Molsidomine) and NO synthesis inhibitor (5 μM DiPhenyleneiodonium) see Subban et al^30^.

### Leaves Preparation and shoot and root Regeneration

Leaves were detached from sterile plants, and the midrib was removed. The leaf blade was cut into segments about 25 mm^2^ (5 mm × 5 mm) and placed on a shoot regeneration (Reg) medium containing standard MS salts as before^30,38^. The medium was supplemented with 30 g l^-1^ sucrose and 8 g l^-1^ agar and the following growth regulators: 4.57 μM IAA; 9.29 μM Kinetin, and 4.56 μM Zeatin (all from Duchefa Co.) or on 1 mg l^-1^ IBA for rooting. The medium was adjusted to pH 5.6. At least 20 leaf segments were placed on each Petri dish with at least four plates per treatment in all experiments. Shoot regeneration from leaf segments was scored 30–32 days after putting them on the medium; root regeneration from segments was scored 10-15 days after placing the leaf segment on medium. Leaf petiole rooting was scored between 4 to 15 days.

Analyses of variance (ANOVA) were performed with the SAS/JMP software (SAS Institute Inc., Cary, NC, USA). Differences among means were calculated based on the Tukey–Kramer honestly significant difference (HSD) test for three or more treatments and T-test for two treatments^30^.

The numbers of explants and regenerated shoots or roots were scored. The regenerative capacity is represented by the number of regenerated shoots or roots per explants. At least three independent experiments (biological triplicates) were performed, and in each, at least three samples were tested.

### RNA Preparation and Transcript Detection

RNA was isolated from leaf segment using the TRI-reagent (Sigma-Aldrich, St. Louis, MO, USA) according to the manufacturer’s instructions and then treated with DNAse (Turbo DNA-free™, Ambion Waltham, Massachusetts, USA). Total RNA was sent to the Weizmann Institute G-INCPM unit for RNA-seq analysis.

### Transcriptome analysis

Raw-reads were subjected to a filtering and cleaning procedure. The Trimmomatic tool^41^ was used to remove Illumina adapters from the reads. Next, the FASTX Toolkit (http://hannonlab.cshl.edu/fastx_toolkit/index.html, version 0.0.13.2) was used to trim read-end nucleotides with quality scores <30, using the FASTQ Quality Trimmer, and to remove reads with less than 70% base pairs with a quality score ≤30 using the FASTQ Quality Filter. Clean reads were mapped to the reference genome of *Nicotiana tabacum*^42^ ftp://ftp.solgenomics.net/genomes/Nicotiana_tabacum/edwards_et_al_2017/assembly/) using STAR software^43^. Gene abundance estimation was performed using Cufflinks^44^ combined with gene annotations from the Sol Genomics Network database (https://solgenomics.net/; ftp://ftp.solgenomics.net/genomes/Nicotiana_tabacum/edwards_et_al_2017/annotation). PCA analysis and Heatmap visualization were performed using R Bioconductor^45^. Gene expression values were computed as FPKM. Differential expression analysis was completed using the DESeq2 R package^46^. Genes that varied from the control more than twofold, with an adjusted *p-value* of no more than 0.05, were considered differentially expressed^47^. Venn diagrams were generated using the online tool at bioinformatics.psb.ugent.be/webtools/Venn/.

KOBAS 3.0 tool^48^ http://kobas.cbi.pku.edu.cn/kobas3/?t=1) was used to detect the statistical enrichment of differential expression genes in the KEGG pathway and Gene Ontology (GO).

## Author Contributions

Conceptualization, M.R.; methodology, M.R., A.D.-F; software, A.D.-F; experimental, Y.K., M.F. data curation, A.D.-F.; Original draft preparation, M.R., A.D.-F.; writing, reviewing and editing, M.R.; visualization, M.R. A.D.-F. All authors have read and agreed to the published version of the manuscript.

## Notes

### Competing Interest Statement

The authors have declared no competing interest.

