## Supplementary material for "Florigen governs shoot regeneration": supplemetal

**Flowering locus T determines shoot regeneration competence.**

Supplemental material

Percent rooting

| Leaf number | % |
| --- | --- |
| 4 | 100 |
| 5 | 100 |
| 6 | 100 |
| 7 | 100 |
| 8 | 71.4286 |
| 9 | 62.5 |
| 10 | 62.5 |
| 11 | 55.5556 |
| 12 | 77.7778 |
| 13 | 60 |
| 14 | 70 |
| 15 | 55.5556 |
| 16 | 62.5 |
| 17 | 16.6667 |
| 18 | 40 |
| 19 | 0 |
| 20 | 0 |
| 21 | 0 |

**Figure S1**: Percent rooting of tobacco leaves. Tobacco plants were grown in pots, and leaves were detached from the stem with the petioles. The leaves petioles were placed in agar containing one mg/L IBA with 2% sucrose. Rooting was calculated as the number of leaves that produced roots from a batch of leaves at the same location on the stem. Original rooting is shown in figure 1.

**c**

**b**

**a**

Percet shooting (%)

| Genotype | % |
| --- | --- |
| Juvenile | 100 |
| Mature | 60.1518 |

Percent rooting (%)

| Days on IBA | Leaf 4 | Leaf 8 | Leaf 20 |
| --- | --- | --- | --- |
| 9 | 0 | 0 | 0 |
| 11 | 12.772 | 20.695 | 1.42857 |
| 13 | 23.3838 | 30.8369 | 1.25 |
| 15 | 42.7622 | 44.6825 | 2.39899 |
| 19 | 44.4143 | 51.4502 | 5.65909 |

Leave at flowering

| Genotype | leaves |
| --- | --- |
| SR1-WT | 17 |
| SR1+florigen-like #10 | 11.25 |
| SR1+florigen-like #11 | 7 |

**Figure S2**: Percent shooting **(a)** or rooting **(b)** of tobacco leaf segments. Tobacco plants were grown in pots, and leaves were detached from the stem and cut into segments. The leaf segments were placed on Reg medium agar containing 3% sucrose for shooting and agar containing one mg/L IBA for rooting. Shooting or Rooting percent was calculated as the number of segments that produced shoots or roots from a batch of leaf segments on the same plate. **c**: Avocado floeigen induces early flowering when expressed in tobacco plants.

width to Length ratio

| Leaf number | Control |
| --- | --- |
| 1 | 1.1216 |
| 2 | 1.0034 |
| 3 | 0.985722 |
| 4 | 0.888195 |
| 5 | 0.787841 |
| 6 | 0.760738 |
| 7 | 0.734236 |
| 8 | 0.806024 |
| 9 | 0.777878 |
| 10 | 0.827963 |
| 11 | 0.814276 |
| 12 | 0.748882 |
| 13 | 0.71885 |
| 14 | 0.792495 |
| 15 | 0.78375 |
| 16 | 0.732383 |
| 17 | 0.692625 |
| 18 | 0.629096 |
| 19 | 0.6238 |
| 20 | 0.583748 |
| 21 | 0.647059 |

| Leaf number | Diphenyl |
| --- | --- |
| 5 | 0.813559 |
| 6 | 0.826446 |
| 7 | 0.811111 |
| 8 | 0.889551 |
| 9 | 0.858491 |
| 10 | 0.818632 |
| 11 | 0.804213 |
| 12 | 0.79393 |
| 13 | 0.75135 |
| 14 | 0.740089 |
| 15 | 0.746576 |
| 16 | 0.737806 |
| 17 | 0.705867 |
| 18 | 0.707899 |
| 19 | 0.747851 |
| 20 | 0.714093 |
| 21 | 0.716577 |

| Leaf number | Molsido |
| --- | --- |
| 7 | 0.791946 |
| 8 | 0.844744 |
| 9 | 0.853393 |
| 10 | 0.841385 |
| 11 | 0.820588 |
| 12 | 0.817031 |
| 13 | 0.831552 |
| 14 | 0.792233 |
| 15 | 0.812675 |
| 16 | 0.783985 |
| 17 | 0.844055 |
| 18 | 0.906814 |
| 19 | 0.761221 |
| 20 | 0.75398 |
| 21 | 0.699774 |
| 22 | 0.700208 |

**Figure S3**: Width to length ratio of tobacco leaves. Tobacco plants were grown in pots and spray at the 7 to 8 leaf stage with either water or Diphenyl (5 μM) or Molsido (5 μM) solutions. The leaves' length and width were measured at flowering, and the ratio width to length was calculated. Each point is the mean±se of at least five individual plants.

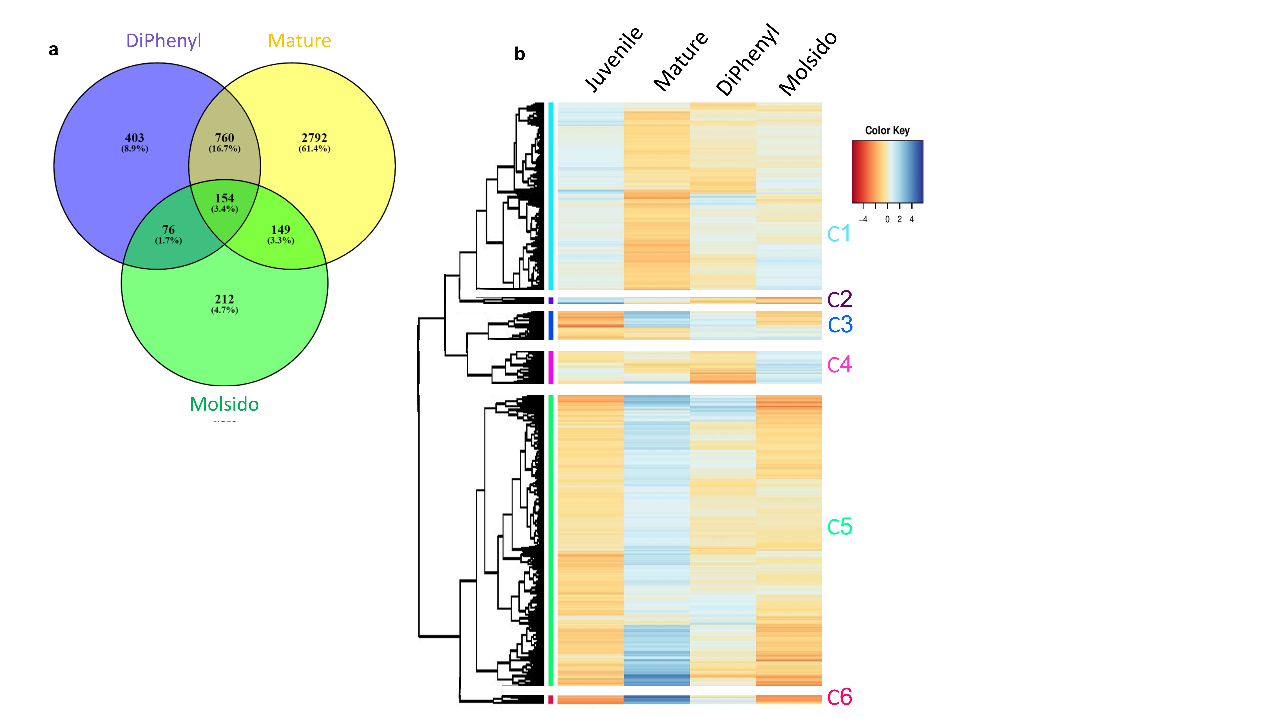

**Fig. S4**: Differential gene expression between juvenile and mature leaves and juvenile leaf treated with DiPhenyl and juvenile leaf treated with Molsido **a**: Number of mRNAs expressed differently than the juvenile samples. Plants grew on an agar medium supplemented with 2% sucrose and 5μM DiPhenyl or 5μM Molsido. At the seven-leaf stage, leaf blade disks were cut and RNA extracted and sent to RNA-seq. The experiment was done in two replicates with five plants each. **b**: Visualisation by heatmap of genes clusters whose mRNA expression increase or decreases at the juvenile, mature, juvenile leaf treated with DiPhenyl and juvenile leaf treated with Molsido.

**Table S1**: Expression level of possible *TEMPRANILLO* gene in tobacco juvenile or mature leaves or juvenile leaves from plants treated with Diphenyl (5 μM) or Molsido (5 μM) after germination.

| Gene ID | Gene description | Juvenile expression level±SD | Juvenile +Diphenyl expression level±SD | Mature expression level±SD | Juvenile +Molsido expression level±SD |
| --- | --- | --- | --- | --- | --- |
| Nitab4.5_0000188g0320 | B3 domain-containing protein At3g18960 IPR003340 Transcriptional factor B3 | 13.0±0.0 | 9.1±0.6 | 12.3±0.0 | 11.8±0.0 |
| Nitab4.5_0000316g0140 | B3 domain-containing protein At1g20600 IPR003340 Transcriptional factor B3 | 0.0±0.0 | 0.0±0.0 | 0.0±0.0 | 0.0±0.0 |
| Nitab4.5_0000380g0010 | B3 domain-containing protein Os03g0212300 IPR003340 Transcriptional factor B3 | 9.2±2.4 | 13.2±0.7 | 12.9±3.8 | 12.2±1.3 |
| Nitab4.5_0000519g0250 | AP2 domain-containing transcription factor IPR003340 Transcriptional factor B3 | 0.0±0.1 | 0.0±0.1 | 0.0±0.0 | 0.0±0.1 |
| Nitab4.5_0000573g0030 | Auxin response factor 24 IPR003340 Transcriptional factor B3 | 6.8±0.8 | 23.1±9.6 | 15.6±2.0 | 6.2±0.3 |
| Nitab4.5_0000733g0200 | B3 domain-containing protein At3g18960 IPR003340 Transcriptional factor B3 | 9.1±1.9 | 8.0±1.2 | 9.8±0.5 | 10.0±0.4 |
| Nitab4.5_0000799g0050 | Auxin response factor 24 IPR003340 Transcriptional factor B3 | 10.4±1.5 | 18.1±6.5 | 25.8±4.3 | 18.7±1.7 |
| Nitab4.5_0000813g0020 | B3 domain-containing transcription factor ABI3 IPR003340 Transcriptional factor B3 | 0.0±0.0 | 0.0±0.0 | 0.0±0.0 | 0.0±0.0 |
| Nitab4.5_0001034g0090 | B3 domain-containing protein Os01g0234100 IPR003340 Transcriptional factor B3 | 6.0±0.2 | 6.1±0.7 | 8.5±0.1 | 7.1±0.3 |
| Nitab4.5_0001101g0130 | B3 domain-containing protein At5g42700 IPR003340 Transcriptional factor B3 | 0.8±0.8 | 0.1±0.1 | 0.4±0.4 | 0.6±0.6 |
| Nitab4.5_0001657g0040 | B3 domain-containing transcription factor ABI3 IPR003340 Transcriptional factor B3 | 0.0±0.0 | 0.0±0.0 | 0.0±0.0 | 0.0±0.0 |
| Nitab4.5_0001663g0030 | B3 domain-containing protein At3g18960 IPR003340 Transcriptional factor B3 | 3.3±0.1 | 2.2±0.6 | 2.8±0.6 | 3.0±0.8 |
| Nitab4.5_0001888g0040 | B3 domain-containing protein At1g20600 IPR003340 Transcriptional factor B3 | 0.0±0.0 | 0.0±0.0 | 0.0±0.0 | 0.0±0.0 |
| Nitab4.5_0002172g0100 | AP2 domain-containing transcription factor IPR003340 Transcriptional factor B3 | 0.0±0.0 | 0.0±0.0 | 0.0±0.0 | 0.0±0.0 |
| Nitab4.5_0003194g0040 | B3 domain-containing protein Os05g0481400 IPR003340 Transcriptional factor B3 | 0.3±0.1 | 0.6±0.4 | 0.7±0.5 | 0.5±0.2 |
| Nitab4.5_0000351g0250 | NtFT1 | 0.0±0.0 | 3.6±1.9 | 2.1±1.2 | 0.0±0.0 |

**Table S2**: Expression level of possible *squamosa promoter binding-like* gene in tobacco juvenile or mature leaves or juvenile leaves from plants treated with Diphenyl (5 μM) or Molsido (5 μM) after germination.

| Gene ID | Gene description | Juvenile expression level±SD | Diphenyl expression level±SD | Mature expression level±SD | Molsido expression level±SD |
| --- | --- | --- | --- | --- | --- |
| Nitab4.5_0000016g0300 | Squamosa promoter-binding protein IPR004333 Transcription factor, SBP-box | 3.4±0.3 | 3.1±0.1 | 4.9±1.2 | 4.6±0.1 |
| Nitab4.5_0000059g0380 | Squamosa promoter binding protein-like 1 IPR004333 Transcription factor, SBP-box | 10.6±2.2 | 8.5±1.2 | 11.5±1.4 | 10.7±0.3 |
| Nitab4.5_0002994g0100 | Squamosa promoter binding protein-like 1 IPR004333 Transcription factor, SBP-box | 38.3±0.9 | 40.5±7.0 | 49.8±3.3 | 31.1±1.9 |
| Nitab4.5_0000222g0290 | Squamosa promoter binding protein-like 1 IPR004333 Transcription factor, SBP-box | 62.6±6.6 | 64.2±14.6 | 82.2±6.6 | 50.0±3.0 |
| Nitab4.5_0000225g0020 | Squamosa promoter binding protein-like 1 IPR004333 Transcription factor, SBP-box | 1.4±0.4 | 0.8±0.4 | 1.7±0.0 | 3.4±0.4 |
| Nitab4.5_0000363g0110 | Squamosa promoter binding protein-like 1 IPR004333 Transcription factor, SBP-box | 10.0±0.8 | 7.4±1.0 | 11.1±0.7 | 12.1±1.2 |
| Nitab4.5_0000700g0020 | Squamosa promoter binding protein-like 1 IPR004333 Transcription factor, SBP-box | 7.4±0.7 | 6.4±0.5 | 7.5±1.6 | 7.5±0.1 |
| Nitab4.5_0000745g0150 | Squamosa promoter binding protein-like 1 IPR004333 Transcription factor, SBP-box | 29.4±2.2 | 18.2±4.1 | 50.1±2.3 | 28.9±2.0 |
| Nitab4.5_0000991g0020 | Squamosa promoter-binding protein IPR004333 Transcription factor, SBP-box | 0.0±0.0 | 0.1±0.1 | 0.6±0.6 | 0.2±0.1 |
| Nitab4.5_0000638g0040 | Squamosa promoter binding-like protein IPR004333 Transcription factor, SBP-box | 1.2±0.1 | 1.1±0.5 | 1.8±0.5 | 2.2±0.3 |
| Nitab4.5_0000210g0190 | Squamosa promoter binding-like protein IPR017238 Squamosa promoter-binding protein | 0.2±0.0 | 0.6±0.4 | 0.8±0.5 | 0.0±0.0 |
| Nitab4.5_0002219g0060 | Squamosa promoter binding-like protein IPR004333 Transcription factor, SBP-box | 3.2±1.0 | 0.6±0.6 | 8.6±2.8 | 6.0±1.0 |
| Nitab4.5_0002299g0030 | Squamosa promoter-binding protein IPR004333 Transcription factor, SBP-box | 3.2±0.1 | 13.7±8.2 | 43.5±6.8 | 5.7±0.4 |
| Nitab4.5_0000210g0160 | Squamosa promoter binding protein 1 IPR004333 Transcription factor, SBP-box | 1.8±0.8 | 5.8±2.3 | 12.8±3.2 | 2.5±0.2 |
| Nitab4.5_0001752g0040 | Squamosa promoter binding-like protein IPR004333 Transcription factor, SBP-box | 1.0±0.1 | 0.5±0.3 | 2.1±0.2 | 1.7±0.2 |
| Nitab4.5_0003348g0050 | Squamosa promoter binding-like protein IPR017238 Squamosa promoter-binding protein | 7.0±0.3 | 12.7±5.5 | 18.8±1.3 | 10.9±2.3 |
| Nitab4.5_0004959g0040 | Squamosa promoter binding-like protein IPR004333 Transcription factor, SBP-box | 0.0±0.0 | 0.2±0.0 | 0.1±0.0 | 0.2±0.2 |
| Nitab4.5_0001315g0110 | Squamosa promoter-binding-like protein 11 IPR004333 Transcription factor, SBP-box | 0.8±0.1 | 0.0±0.0 | 0.1±0.1 | 0.7±0.1 |
| Nitab4.5_0000067g0130 | Squamosa promoter-binding-like protein 11 IPR004333 Transcription factor, SBP-box | 24.6±4.7 | 20.0±0.4 | 12.7±3.2 | 27.5±3.5 |
| Nitab4.5_0000861g0050 | Squamosa promoter-binding-like protein 11 IPR004333 Transcription factor, SBP-box | 7.1±2.3 | 1.7±0.1 | 5.7±1.3 | 9.0±0.4 |
| Nitab4.5_0003900g0020 | Squamosa promoter-binding-like protein 11 IPR004333 Transcription factor, SBP-box | 5.0±0.5 | 3.9±0.6 | 7.4±1.6 | 8.7±2.2 |
| Nitab4.5_0000027g0060 | Squamosa promoter binding protein 3 IPR004333 Transcription factor, SBP-box | 4.7±1.0 | 5.9±1.8 | 19.2±3.6 | 5.4±0.7 |
| Nitab4.5_0000509g0030 | Squamosa promoter binding protein 3 IPR004333 Transcription factor, SBP-box | 3.0±1.2 | 1.5±0.6 | 1.0±0.1 | 0.6±0.1 |
| Nitab4.5_0000509g0040 | Squamosa promoter binding protein 3 IPR004333 Transcription factor, SBP-box | 1.6±0.3 | 0.8±0.8 | 0.7±0.1 | 0.8±0.2 |
| Nitab4.5_0001118g0090 | Squamosa promoter binding protein 3 IPR004333 Transcription factor, SBP-box | 0.8±0.3 | 0.8±0.1 | 0.5±0.2 | 0.7±0.2 |
| Nitab4.5_0001328g0090 | Squamosa promoter binding protein 3 IPR004333 Transcription factor, SBP-box | 6.2±0.2 | 4.5±0.8 | 1.3±0.3 | 4.4±0.6 |
| Nitab4.5_0001328g0100 | Squamosa promoter binding protein 3 IPR004333 Transcription factor, SBP-box | 1.2±0.6 | 0.8±0.6 | 0.8±0.4 | 2.1±0.1 |
| Nitab4.5_0001797g0070 | Squamosa promoter binding protein 3 IPR004333 Transcription factor, SBP-box | 4.7±1.7 | 4.7±3.1 | 1.4±0.3 | 9.8±1.6 |
| Nitab4.5_0002467g0020 | Squamosa promoter binding protein 3 IPR004333 Transcription factor, SBP-box | 4.3±1.5 | 5.8±1.4 | 13.1±0.5 | 8.0±1.0 |
| Nitab4.5_0000351g0250 | NtFT1 | 0.0±0.0 | 3.6±1.9 | 2.1±1.2 | 0.0±0.0 |
